# The role of bacterial size, shape and surface in macrophage engulfment of uropathogenic *E. coli* cells

**DOI:** 10.1101/2022.11.20.517300

**Authors:** Elizabeth Peterson, Bill Söderström, Nienke Prins, Giang H.B. Le, Lauren E. Hartley-Tassell, Chris Evenhuis, Rasmus Birkholm Grønnemose, Thomas Emil Andersen, Jakob Møller-Jensen, Gregory Iosifidis, Iain G. Duggin, Bernadette Saunders, Elizabeth J. Harry, Amy L. Bottomley

## Abstract

Uropathogenic *Escherichia coli* (UPEC) can undergo extensive filamentation in the host during acute urinary tract infections (UTIs). It has been hypothesised that this morphological plasticity allows bacteria to avoid host immune responses such as macrophage engulfment. However, it is still unclear what properties of filaments are important in macrophage-bacteria interactions. The aim of this work was to investigate the contribution of bacterial biophysical parameters, such as cell size and shape, and physiological parameters, such as cell surface and the environment, to macrophage engulfment efficiency. Viable, reversible filaments of known lengths and volumes were produced in the UPEC strain UTI89 using a variety of methods, including exposure to cell-wall targeting antibiotics, genetic manipulation and isolation from an *in vitro* human bladder cell model. Quantification of the engulfment ability of macrophages using gentamicin-protection assays and fluorescence microscopy demonstrated that the ability of filaments to avoid macrophage engulfment is dependent on a combination of size (length and volume), shape, surface and external environmental factors. UTI89 filamentation was also found to occur independently of the SOS-inducible filamentation genes, *sulA* and *ymfM*, demonstrating the non-essential requirement of these genes for UTI89 filamentation and their ability to avoid macrophage engulfment. With several strains of UPEC now resistant to current antibiotics, our work identifies the importance of bacterial morphology during infection and may provide new ways to prevent or treat these infections via immune modulation or antimicrobials.

**Author Summary:** Urinary tract infections (UTIs) are one of the most common bacterial infections worldwide with 50% of women suffering from a UTI during their lifetime. Escherichia coli is the primary bacteria responsible for UTIs and is usually found in short rod forms. However, during UTIs E. coli can elongate into extremely long thin shapes called ‘filaments’. Filaments are thought to be advantageous during infections because they are too long to be engulfed and killed by immune cells called macrophages. Due to increasing antibiotic resistance in bacteria there is a strong need for the discovery of new ways to treat infections and this is only possible once we thoroughly understand the mechanisms bacteria employ to overcome our immune response. Therefore, we investigated the effect of E. coli filamentation on macrophage engulfment along with other aspects of bacteria reported to influence engulfment. We found that the ability of filaments to avoid macrophage engulfment is dependent on a combination of size (length and volume), shape, surface and external environmental factors. Our research has highlighted the importance of bacterial shape changes during infections and provided a foundational understanding of macrophage engulfment of filaments. Eventually, this knowledge may reveal new targets for treatment of infections.

## Introduction

The phenomenon of rod-shaped bacteria to change shape into long “spaghetti-like” filamentous cells has been repeatedly observed in a variety of species [1, 2] under many different environmental conditions [3]. Filaments arise when cell division is inhibited whilst DNA replication and chromosome segregation continue, with filaments containing multiple copies of the chromosome along their lengths [2, 4, 5]. Whilst this morphology has been commonly observed, the biological impact of filamentation is still largely unknown, although it has been suggested to provide survival advantages in various niches, such as protection against aquatic protist predators for *Flectobacillus* species [6], host intracellular survival for *Burkholderia pseudomallei* [7] or protection against phagocytosis by host immune cells during infection [8, 9]. Understanding why bacteria filament in certain environments will provide insights into their pathogenicity and the potential of this morphology to impede the treatment of infections, particularly those caused by antibiotic resistant bacteria [4, 10–12].

Urinary tract infections (UTIs) are extremely common, affecting around 150 million people globally every year, with uropathogenic *Escherichia coli* (UPEC) causing over 80% of these infections [13, 14]. The only treatments available for UTIs are antibiotics and with several UPEC strains currently resistant to many antibiotics, they are an increasing threat to human health [15]. UPEC can undergo extensive filamentation during acute UTIs as part of the infection cycle [16, 17] though the exact role of this morphological change during the infection cycle is not conclusive [16, 18]. Compared to their rod counterparts, filaments may more strongly adhere to vulnerable host cells, and consequently be more effective at establishing infection [19], and/or be better able to avoid phagocytosis by immune cells and thus subvert host immunity during infection [8, 9]. Identifying new treatment approaches should be possible with a detailed understanding of the role(s) that bacterial morphology of pathogenic bacteria, such as UPEC, in interacting with the host immune system during infection.

UPEC filaments are not efficiently engulfed and killed by neutrophils and macrophages in a mouse model of acute UTI [8, 11, 20]. It has been hypothesised that the increase in bacterial cell size prevents immune cells effectively engulfing and clearing the infection [21, 22]. However, immune cells can engulf large particles, such as apoptotic or cancerous tissue cells, clumps of bacteria or inorganic foreign bodies up to 20 μm long [23, 24], leading to the proposal that it is more than just cell size that results in filaments being engulfed less readily than shorter, rod-shaped cells [20]. Computational modelling and biophysical studies using silica or polystyrene nanoparticle beads have examined the interdependence of shape and size in phagocytosis efficiency, showing both aspect ratio and curvature contribute to the ability of immune cells to efficiently phagocytose; prolate particles are more difficult to engulf because of their high aspect ratios and rounded tips [24, 25]. However, how this interplay between size and shape to prevent immune cell engulfment translates to changes in bacterial morphology in live pathogenic bacteria is still relatively unknown.

Understanding the role of filamentation in avoiding immune cell engulfment, and what biophysical aspects contribute to this avoidance, is further complicated by the significant variation in experimental methods used to generate filaments. These variations include stimuli to induce filamentation, cell viability, bacterial species, use of inert particles and immune models, making it difficult to draw definitive conclusions about what aspect(s) of bacterial filamentation provides a survival advantage [22, 26, 27]. Even the fundamental definition of what constitutes a filament is not consistent, with ‘filaments’ varying from 5 μm to over 200 μm depending on the induction method, and the definition of filament length is inconsistently reported between studies [9, 16, 17, 28]. A complete understanding of the role and significance of filamentation, in the context of UPEC and the implications for UTIs, requires a rigorous exploration of what properties of filaments are important in affecting macrophage engulfment.

Here, we have used controlled conditions to generate UPEC filaments of defined length, allowing us to unravel the importance of biophysical parameters such as cell size and shape, and physiological parameters such as cell surface and the environment, during engulfment by human macrophages. Using gentamicin-protection assays and fluorescence microscopy to quantify the engulfment ability of macrophages, we found that UPEC filaments derived from all conditions tested were engulfed less by macrophages compared to their rod counterparts. There was an interplay between size and shape of cells, where spherical cells were more readily engulfed compared to both rods and filaments. Mannose-dependent interaction between UPEC and human macrophages was confirmed as the major pathway for engulfment, regardless of bacterial cell size. Surprisingly, deletion of the widely-reported mannose-binding fimbrial gene *fimH* increased engulfment of both rods and filaments and abolished the size dependency, raising the possibility that there are also mannose-independent interactions between UPEC and macrophages that are critical for engulfment. Decreased complete engulfment of UPEC filaments isolated from an *in vitro* bladder model was also observed. However, an increase in partial engulfment of bladder model filaments compared to antibiotic-induced filaments demonstrated an influence of the bacterial growth environment on engulfment effectiveness. Finally, deletion of the SOS response genes *sulA* and *ymfM* had no effect on the ability of UPEC to filament or influence UPEC engulfment by macrophages. We conclude from these observations that while the size and shape of UPEC are major influencers of effective macrophage engulfment, the surface and growth environment of UPEC rods and filaments also play roles in macrophage engulfment.

## Results

### Cephalexin treatment results in a population of viable UTI89 filaments of defined length

We initially produced viable *E. coli* UTI89 filaments of defined length using cephalexin since it targets PBP3/FtsI to block cell division [29, 30] and is used in the treatment of UTIs [31, 32]. The minimum inhibitory concentration (MIC) of cephalexin in UTI89 was determined to be 5 μg/ml (data not shown) and, to induce filamentation, antibiotic concentrations used were the same as the range detected in the plasma of healthy individuals who have undergone antibiotic treatment (7.7-12.3 μg/ml [32]). At 10 μg/ml, cephalexin-induced filaments had a mean cell length of 10 μm (± 0.3 SEM) and were 6-fold longer than untreated rods (1.7 μm ± 0.02 SEM; Fig 1A, Fig S1A-B). To control for the effect of antibiotic exposure which may affect macrophage engulfment but is unrelated to cell length changes, UTI89 was exposed to a low (sub-MIC) concentration of cephalexin (2.5 μg/ml). These cells had a mean cell length of 2.4 μm (± 0.06 SEM; Fig 1A) and were not significantly different in length to untreated rods. Bacteria from all three populations were viable as measured by membrane permeability and ATP level assays (Fig S2A and S3A). As with UPEC filamentation during the infection cycle [18], cephalexin-induced filamentation was reversible; removal of cephalexin resulted in the resumption of cell division, within 1 hour, producing an almost uniform population of rods by 2 hours incubation in LB media (Fig S4A and S4D). Therefore, for the purpose of this study, we defined filaments as cells that were over 4 μm in length and rods were defined as less than 4 μm long.

**Fig 1.**
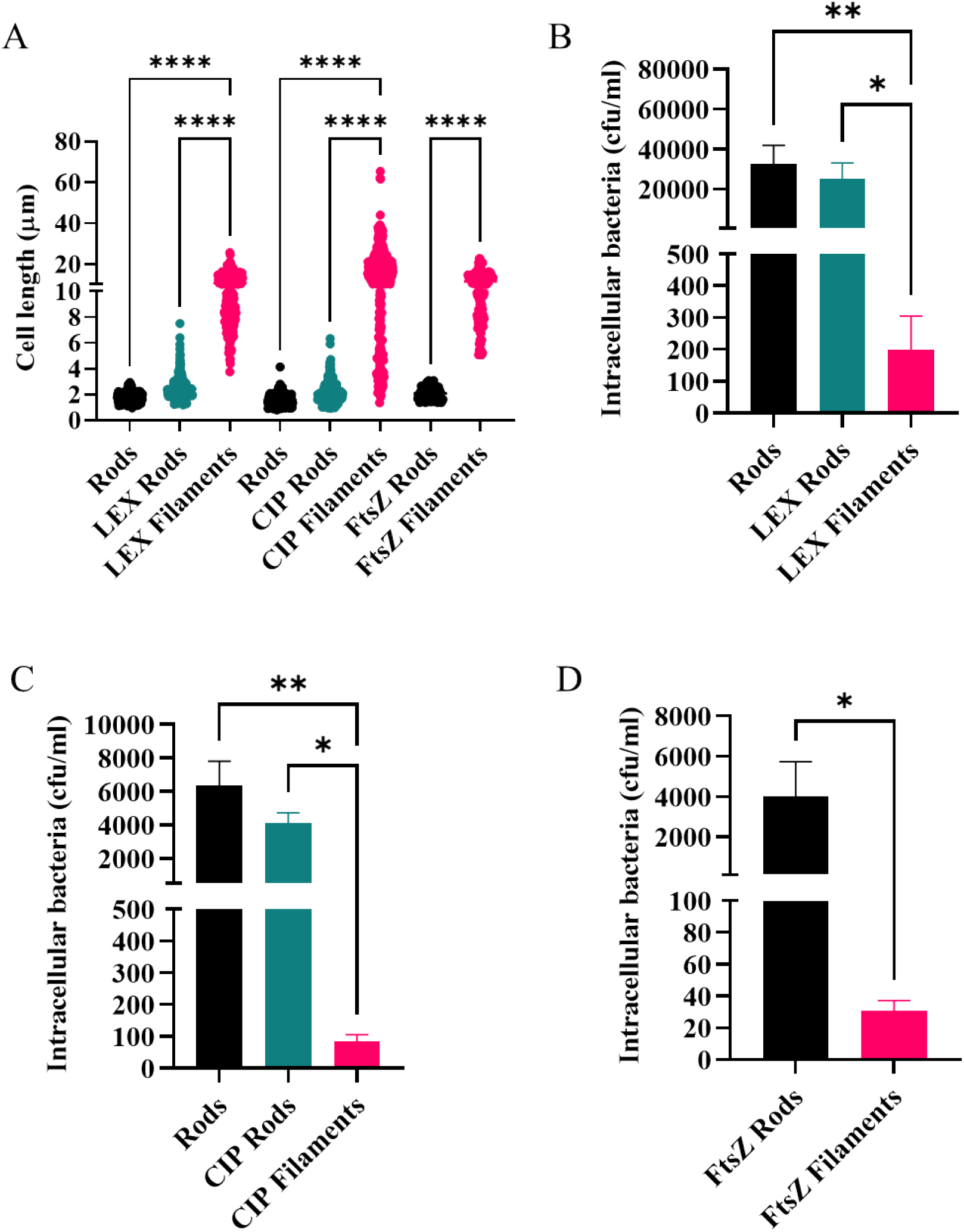
THP-1 macrophages engulf UTI89 filaments significantly less than rods. *UTI89 was treated with either cephalexin (rods: untreated, LEX rods: 2.5 μg/ml, LEX filaments: 10 μg/ml) or ciprofloxacin (rods: untreated, CIP rods: 3.75 ng/ml, CIP filaments: 15 ng/ml). UTI89/pLau80 was treated with 0.2% glucose (FtsZ rods) or 0.2% arabinose to induce expression of* ftsZ-yfp *(FtsZ filaments). (A) Bacterial lengths as determined by phase contrast microscopy with n = 110-426. (B-D) THP-1 macrophages were infected (MOI 10) for 1 hour and intracellular bacterial loads were assessed 2-hours post infection in a gentamicin-protection assay. Data are the averages of 4 independent experiments with error bars representing the SEM. * indicates* p *<0.05*, ** p <*0.01, \*\*\*\** p <*0.0001, determined by one-way ANOVA with multiple comparisons*.

### Cephalexin- and ciprofloxacin-induced filamentation of UTI89 protects against macrophage engulfment

The THP-1 human monocyte cell line (ATCC^®^ TIB-202™), differentiated with phorbol-12-myristate-13-acetate (PMA) to form macrophages, was used to model phagocytic engulfment of cephalexin-induced UTI89 filaments and untreated rods in a gentamicin-protection assay. A gentamicin-protection assay with an infection time of 1 hour and a gentamicin kill time of 1 hour was used in this study following previously published methods [33], and 1 hour was determined to be the maximum amount of time macrophages could be incubated with filaments before the filament population reverted to a rod population (S5 Fig). A multiplicity of infection (MOI) of 10 was used as for other UPEC engulfment studies [33–35]. We established that cephalexin-induced filaments do not revert to rods in the cell assay growth media within the time frame of the assay (Fig S5).

To be able to directly compare the CFU/ml values for rods and filaments given that they differ in their mass, we examined the relationship between CFU/ml and absorbance (bacterial biomass) to ensure that one filament, no matter what length, resulted in one CFU. Cephalexin treatment resulted in filaments with a mean length 6.1-fold that of rods. We found, by measuring the absorbance and CFU/ml of these populations over time, that the biomass of populations increases steadily for all populations (Fig S6A) but after the addition of cephalexin there is little increase in CFU/ml of filaments (Fig S6B). This implies that while cell division to form more cells stops for filaments after cephalexin addition, length and therefore biomass continues to increase. Furthermore, the CFU/ml of rods is approximately 6-fold greater than that of filaments (Fig S6B), demonstrating a clear relationship with the 6.1-fold length increase in filament length over the same time. From this data we can conclude that 1 colony forming unit (cfu) will approximately equal 1 bacterium (rod or filament), however filaments could account for over 6 rods in biomass. This validates our comparisons of rod and filament engulfment measured by CFU/ml in gentamycin-protection assays.

Gentamicin-protection assays revealed that cephalexin-induced filaments (LEX filaments) were engulfed by macrophages 150-fold less frequently than untreated rods (Fig 1B; p = 0.009). There was no significant change to macrophage engulfment levels between low dose cephalexin-exposed rods (2.5 μg/ml; LEX rods) and untreated cells (Fig 1B), suggesting that the reduction of ability of macrophages to engulf filaments is at least in part a consequence of the morphological change of filamentation caused by antibiotic treatment, rather than just antibiotic exposure *per se* (see also FtsZ-YFP expression induced-filaments below).

We next examined whether induction of UTI89 filamentation, using an antibiotic that has a different target to PBP3 but is also used in the treatment of UTIs [36], leads to a similar decrease in macrophage engulfment. Cells treated with 15 ng/ml ciprofloxacin, which targets DNA gyrase [37], resulted in viable, reversible filamentation in which cells were 10-fold longer than both untreated and low-dose ciprofloxacin-treated (3.75 ng/ml) control cells (Fig 1A, Fig S1C-D, Fig S2B, Fig S3B, Fig S4B). As with cephalexin-induced filamentation, long cells produced by ciprofloxacin treatment of UTI89 (CIP filaments) provided significant protection against macrophage engulfment; being engulfed 60-fold less than their rod counterpart (Fig 1C). As observed for LEX rods, low-dose ciprofloxacin treated cells (CIP rods) were similarly engulfed compared to untreated UTI89 (Fig 1C). Overall, these results indicate that filamentation provides an advantage during macrophage engulfment, regardless of the antibiotic stimuli to induce filamentation.

### UTI89 filamentation induced by abnormal FtsZ expression also protects against engulfment by macrophages

We found that exposure to low doses of antibiotic did not alter macrophage engulfment compared to untreated cells, suggesting that it is increased cell length, and not antibiotic treatment, that results in drastically low engulfment of filaments. To confirm this, we used a stimulus that did not rely on antibiotics to induce filamentation. FtsZ was produced (as a YFP fusion) from pLau80 [38] under P_BAD_ control, in addition to endogenous FtsZ in UTI89. This induced strong filamentation [39] on addition of arabinose (herein referred to as FtsZ filaments) [40]. FtsZ filaments were 5.5-fold longer than their rod counterpart (11.4 μm ± 0.4 SEM vs. FtsZ rods; 2.1 μm ± 0.04 SEM; Fig 1A; Fig S1E-F) and were as viable as FtsZ rods (Fig S2C) but showed a moderate decrease (29%, *p* = 0.002) in ATP production compared to FtsZ rods (Fig S3C). This could be due to the overabundance of FtsZ (as a combination of FtsZ and FtsZ-YFP) affecting metabolism when FtsZ levels are usually tightly controlled [41]. As with the antibiotic-induced filaments, filamentation was reversible once *ftsZ-yfp* expression was repressed (Fig S4C), albeit taking a longer time to revert to rods than the removal of antibiotics (by 2 hours over 50% of cells remained over 4 μm in length), presumably due to the need to reduce the level of FtsZ-YFP appropriately. Macrophage engulfment of FtsZ filaments was drastically decreased (100-fold) compared to FtsZ rods (Fig 1D), showing that non-antibiotic induced filamentation also provides substantial protection against macrophage engulfment. The combined results above demonstrate that the actual length of the UTI89 filament is at least one key factor in providing this protection.

### Rods are preferentially engulfed by macrophages in mixed populations of rods and filaments

We have shown that upon various stimuli, viable filaments are engulfed by macrophages significantly less than rods. However, this is for homogenous rod or filament populations. A mixed population of rods and filaments is the norm in most environments [17, 42, 43] which may influence the engulfment dynamics of macrophages. We therefore investigated macrophage engulfment of a mixed rod and filament UTI89 population using cephalexin to stimulate filament formation since this treatment resulted in a uniform mean filament length. We used strains UTI89 and UTI89/pGI5 (the latter constitutively producing monomeric superfolder GFP, msfGFP, from pGI5 [44]) to produce separate populations of viable rods and msfGFP-labelled filaments which were subsequently combined to produce a mixed population of rods (1.6 μm ± 0.03 SEM) and filaments (10.3 μm ± 0.2 SEM). A UTI bladder infection model is comprised of approximately 80% UTI89 filaments (measured via microscopy)[45], and to mimic this, we created a heterogeneous population of approximately 76% filaments and 24% rods, based on known CFU/ml data (Fig 2A). Using a gentamicin-protection assay with the mixed population of UTI89 and UTI89/pGI5 (msfGFP) cells, macrophage engulfment was 10-fold less than a UTI89 rod-only population (Fig 2B), indicating reduced overall engulfment in a heterogeneous population. However, while only 24% of the mixed population added to macrophages was rods, 76% of the plate-recovered engulfed bacteria consisted of rods as determined by selective antibiotic plating (due to spectinomycin resistance from pGI5 (msfGFP); Fig 2A and B), indicating that there is also a distinct preference for engulfment of rods over filaments in a heterogeneous mixed population. The plasmid had no effect on engulfment, and it made no difference which population (rods or filaments) contained the plasmid (data not shown).

**Fig 2.**
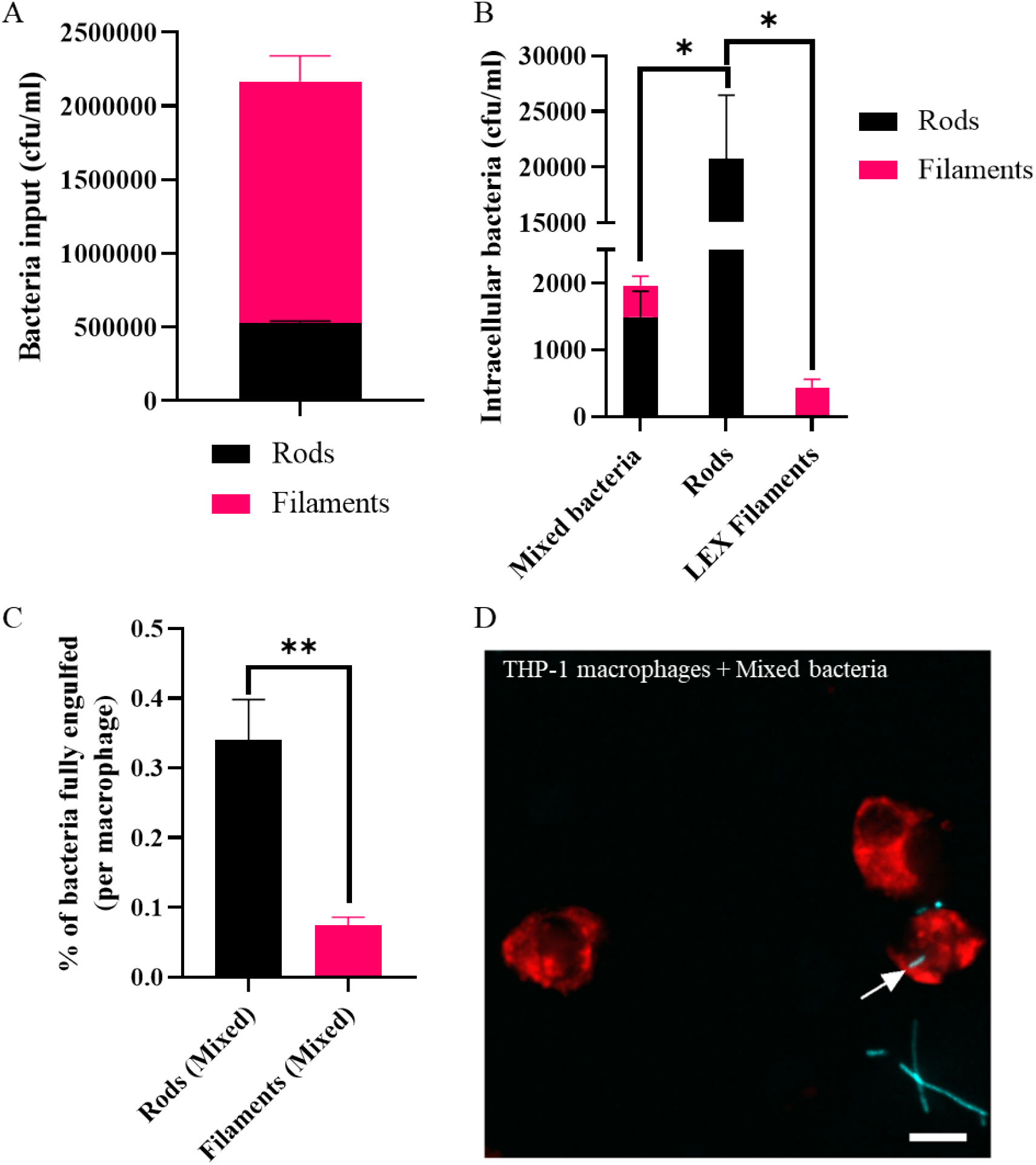
In a heterogeneous population of rods and filaments, rods are engulfed by THP-1 macrophages preferentially over filaments. *UTI89 and UTI89/pGI5 (msfGFP) were untreated (rods) or exposed to 10 μg/ml cephalexin (LEX filaments). (A) rods and filaments were combined to create one population (Mixed bacteria) of 24% rods and 76% filaments. (B) THP-1 macrophages were infected (MOI 10) for 1 hour and intracellular bacterial loads were assessed 2-hours post infection in a gentamicin-protection assay. (C-D) Macrophages were infected with UTI89/pGI5 (msfGFP) (cyan) Mixed bacteria at MOI 10, fixed after 60 minutes and stained with 1X CellMask™ Orange (red). Images were acquired using a DeltaVision Elite microscope with the 40X dry NA 0.60 objective. (C) Percentage of bacterial populations fully engulfed (number of bacteria fully engulfed/total number of bacteria counted X 100) is normalised and presented as per macrophage to allow for comparisons despite variations in number of macrophages counted from microscopy images. (D) Representative image (maximum intensity projection) of infected macrophage with arrow indicating an internalised rod bacterium. Scale bar = 10 μm. Data are the averages of 4 independent experiments (A and B) or 2 independent experiments (C) with error bars representing the SEM. * indicates* p < *0.05*, ** p < *0.01, determined by one-way ANOVA with multiple comparisons or Welch’s t-test*.

To directly visualize the interaction between THP-1 macrophages and bacteria, widefield fluorescence microscopy was performed following a 60-minute incubation with a mixed population of UTI89/pGI5 (msfGFP) rods and filaments. Cell lengths of bacteria were measured and showed that rods (bacteria < 4 μm in length, to account for cells preparing to divide) were engulfed preferentially over filaments (bacteria > 4 μm in length) after a 1-hour infection (Fig 2C-D). Microscopy analysis of fully engulfed bacteria (bacteria completely enveloped by the macrophage membrane) showed that in the artificial mixed population the rod populations were fully engulfed significantly more than filaments. Approximately 0.35% of the rod populations added were fully engulfed by each macrophage, compared to less than 0.1% of the filament population, showing a 3.5-fold difference in successful engulfment (Fig 2C). Therefore, our data shows that as with separate populations, rods are engulfed preferentially over filaments in a mixed population.

### As cell length increases macrophage engulfment decreases at an exponential rate

Previous studies, primarily examining non-viable particles, have demonstrated that the optimal particle size for engulfment by macrophages is between 1 to 3 μm [26, 46–48]. However, macrophages have been reported to engulf very large particles, such as latex beads or apoptotic cells over 20 μm in diameter [27, 49]. We therefore sought to establish how long a bacterial cell needs to be before there is an observable reduction in macrophage engulfment. Cell populations of UTI89 with increasing average lengths were produced by staggering the time of addition of cephalexin (10 μg/ml) to produce cell populations varying in length from 1.7 to 13.3 μm (Fig 3A). Using a gentamicin-protection assay, we found that just doubling the cell length i.e., blocking the first division, from 1.7 μm for untreated cells to 4.2 μm with cephalexin exposure, is enough to decrease macrophage engulfment 5-fold (*p* = 0.039) (Fig 3A). This was surprising since it has been observed that UPEC filaments can get extremely long [18]. Additionally, we found that increasing cell length resulted in an exponential decrease in engulfment, (nonlinear regression R squared value = 0.79, Fig 3B). Similar trends were observed with filaments of varying lengths (between 2 and 8.7 μm) as a result of *ftsZ* expression (using UTI89/pLau80/pGI5 (*ftsZ-yfp*, msfGFP) cultures) (Fig 3C and 3D). Overall, while only a small increase in cell length (2.4-fold) is enough to cause a significant reduction in macrophage engulfment, continued increase in cell length results in a corresponding reduction in engulfment efficiency. Thus, there does not appear to be a length threshold that bacteria reach which suddenly decreases engulfment. A relatively small increase in cell length is enough to reduce macrophage engulfment and as length increases, engulfment continues to decrease at an exponential rate.

**Fig 3.**
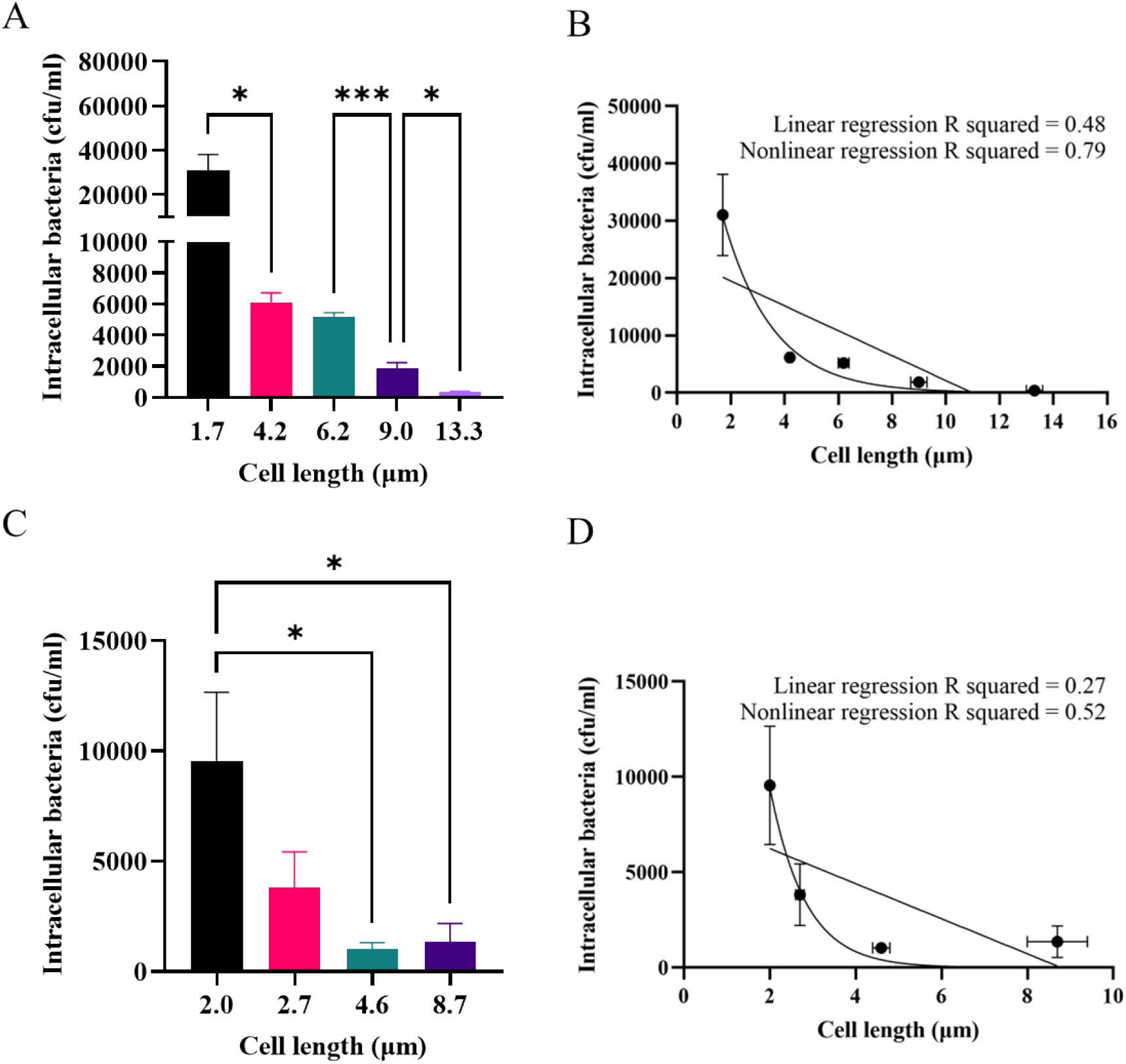
THP-1 macrophage engulfment of UTI89 decreases exponentially as bacterial length increases. (*A-B) UTI89 was treated with 10 μg/ml cephalexin for increasing periods of time to produce filaments of increasing average length. (C-D) UTI89/pLau80 was treated with 0.2% (v/v) arabinose for increasing periods of time to produce FtsZ filaments of increasing average length. (A-D) THP-1 macrophages were infected (MOI 10) for 1 hour and intracellular bacterial loads were assessed 2-hours post infection in a gentamicin-protection assay. Data are the averages of 4 independent experiments with vertical error bars representing the SEM of intracellular bacteria and horizontal error bars (B and D) represent SEM of bacterial length, n = 105-250. * indicates* p < *0.05*, ** p < *0.01*, *** p < *0.001, determined by one-way ANOVA with multiple comparisons. R squared values generated from linear and nonlinear regression analyses, which were plotted as straight or curved lines, respectively, on B and D*.

### Macrophage engulfment is influenced by both shape and size of UTI89 cells

Previous studies using small inert polystyrene particles have indicated that it is the target shape rather than size that influences macrophage engulfment [26, 50]. To establish whether this holds true for live bacterial cells, we created distinct shapes of UTI89 cells using two cell wall-targeting antibiotics. Cephalexin targets PBP3 (FtsI) to inhibit septal cell wall synthesis, resulting in filamentation [30, 51], whilst mecillinam targets PBP2 to inhibit lateral cell wall synthesis, resulting in spherical cells [52–54]. By varying the time of exposure to individual antibiotics, or by combining treatments, cells with different sizes and shapes could be produced (Fig 4A). Cell volumes were measured to be able to compare the size of cells of different shapes.

**Fig 4.**
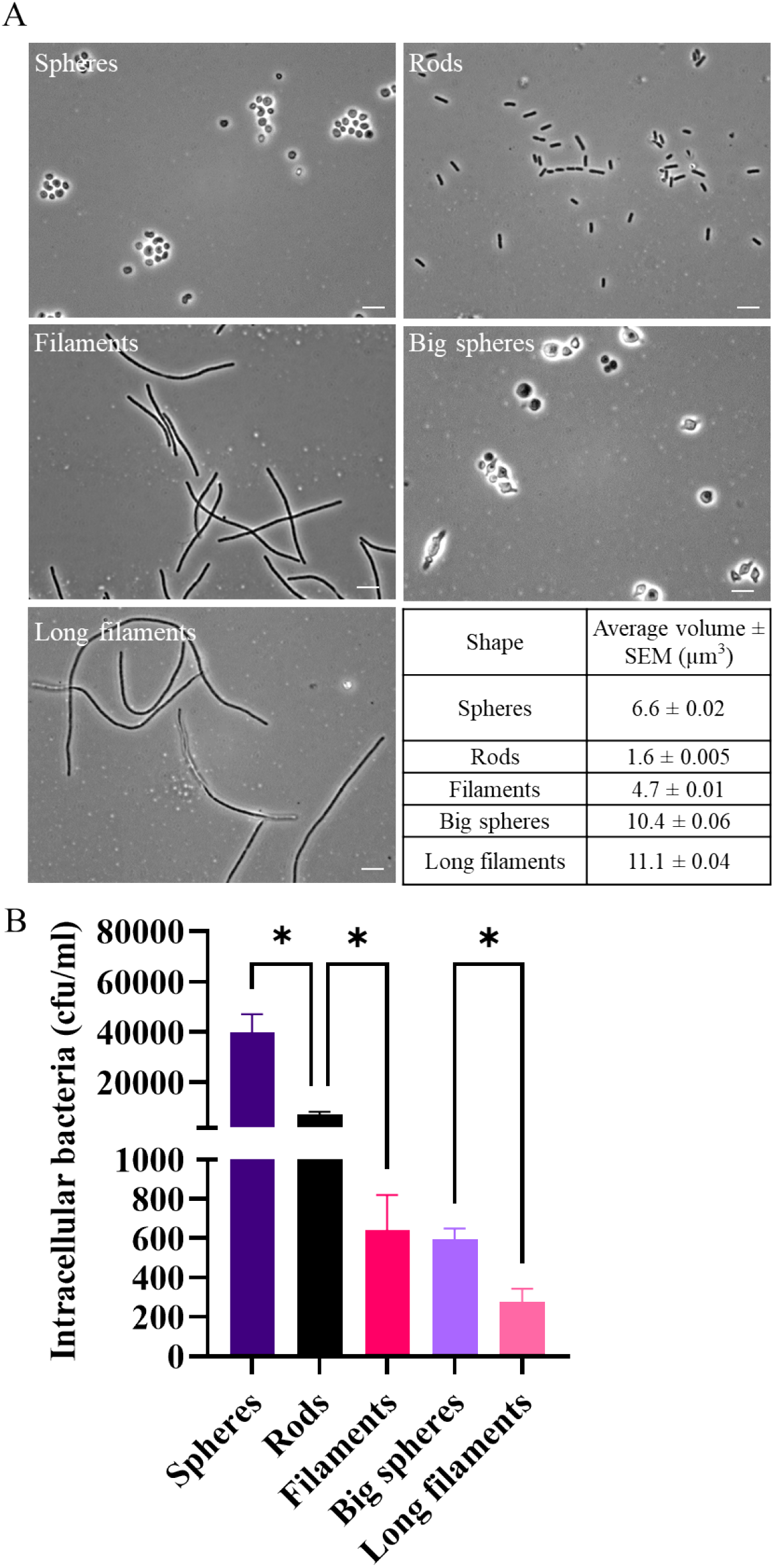
THP-1 macrophages have bacterial size and shape preferences during engulfment. *UTI89 cells were untreated (rods) or treated with 10 μg/ml cephalexin (filaments), 10 μg/ml mecillinam (Spheres), and grown for 3 additional hours with 10 μg/ml mecillinam and cephalexin (Big spheres) or with 10 μg/ml cephalexin (Long filaments). (A) Phase contrast images were acquired using a Zeiss Axioplan 2 microscope with the 100X oil immersion NA 1.4 objective. Images are representative from 1 experiment. Scale bar = 5 μm. Table of average cell volumes with SEM as measured by a Coulter counter, n = 36927-281142. (B) THP-1 macrophages were infected (MOI 10) for 1 hour and intracellular bacterial loads were assessed 2 hours post infection in a gentamicin-protection assay. Data are the averages of 3 independent experiments with error bars representing the SEM. * indicates* p < *0.05, determined by one-way ANOVA with multiple comparisons*.

Although untreated rods had the smallest mean cell volume of 1.6 μm^3^, we found that spherical mecillinam-treated UTI89 with a mean cell volume of 6.6 μm^3^, were engulfed 4-fold more than rods (Fig 4B), indicating that there is primarily a shape preference during engulfment. This is further demonstrated by comparing cells of different shape but similar volume: Spheres (6.6 μm^3^) were engulfed 62-fold more than filaments (4.7 μm^3^; Figure 4B). This shape dependency is conserved when cell volume is increased; Big spheres (10.3 μm^3^), produced by treatment with both antibiotics, were engulfed 2.5-fold more than Long filaments (11.1 μm^3^, Fig 4B). However, there is also a size dependency for engulfment, as increasing the size of spherical cells to Big spheres (10.3 μm^3^) decreased engulfment 80-fold compared to Spheres (6.6 μm^3^). This indicates that there is an interplay between size and shape of UTI89 cells that influences the effectiveness of macrophage engulfment.

### The reduced macrophage engulfment of filaments is unlikely to be due to changes in their ability to bind mannose

Since the bacterial cell surface is the initial point of recognition between a macrophage and its target [55], we investigated if surface glycans differed between cephalexin-induced rods and filaments which could account for differences in engulfment efficiency. Glycan microarray analysis revealed differences in the ability of rods, LEX rods and LEX filaments to bind to different glycan moieties (Table S1). However, these differences did not appear to involve glycans known to be involved in influencing macrophage engulfment and are not predicted to be a major contributing factor in reduced engulfment of filaments [56].

A well-characterised primary interaction between UPEC and macrophages is via mannosylated glycoproteins on the macrophage surface [57]. This binding can be competitively inhibited by methyl α-D-mannopyranoside (α-D-MP) [58], which binds to bacterial surface structures, thus blocking or reducing macrophage binding. Indeed it has previously been shown that addition of α-D-MP results in reduced mouse macrophage engulfment of *E. coli* J96 rods [58], demonstrating that engulfment of UPEC is mannose dependent. It is possible that filament engulfment is less efficient than that of rods due to reduced binding of filaments to host mannosylated receptors. If this were the case, then blocking mannose-binding would not lead to a significant decrease in filament engulfment but would decrease rod engulfment. We therefore tested this by adding α-D-MP to UTI89 rods and cephalexin-induced filaments (~10 μm length) and performing a gentamicin-protection assay with THP-1 macrophages. Consistent with previous results [58], engulfment of rods treated with α-D-MP decreased 16-fold compared to the untreated rod population in the presence of α-D-MP (*p* = 0.041; Fig 5). We observed a similar fold-decrease (13-fold, *p* = 0.009) in engulfment of filaments in the presence of α-D-MP relative to untreated filaments (Fig 5). Rods +α-D-MP were still engulfed significantly more (*p* = 0.038) than filaments +α-D-MP demonstrating that the preference for rods remained. The equal proportional decrease in engulfment of both rods and filaments when mannose binding is inhibited indicates that under these conditions the vast difference in THP-1 engulfment of filaments compared to rods is unlikely to be due to changes in their mannose-specific binding. It is also interesting to note that whilst engulfment of rods and filaments is significantly reduced, there is still a proportion of bacteria that are successfully engulfed when mannosylated glycoproteins are competitively inhibited (rods and filaments +α-D-MP), suggesting that there are mannose-independent interactions between UPEC and macrophages that are critical for optimal engulfment.

**Fig 5.**
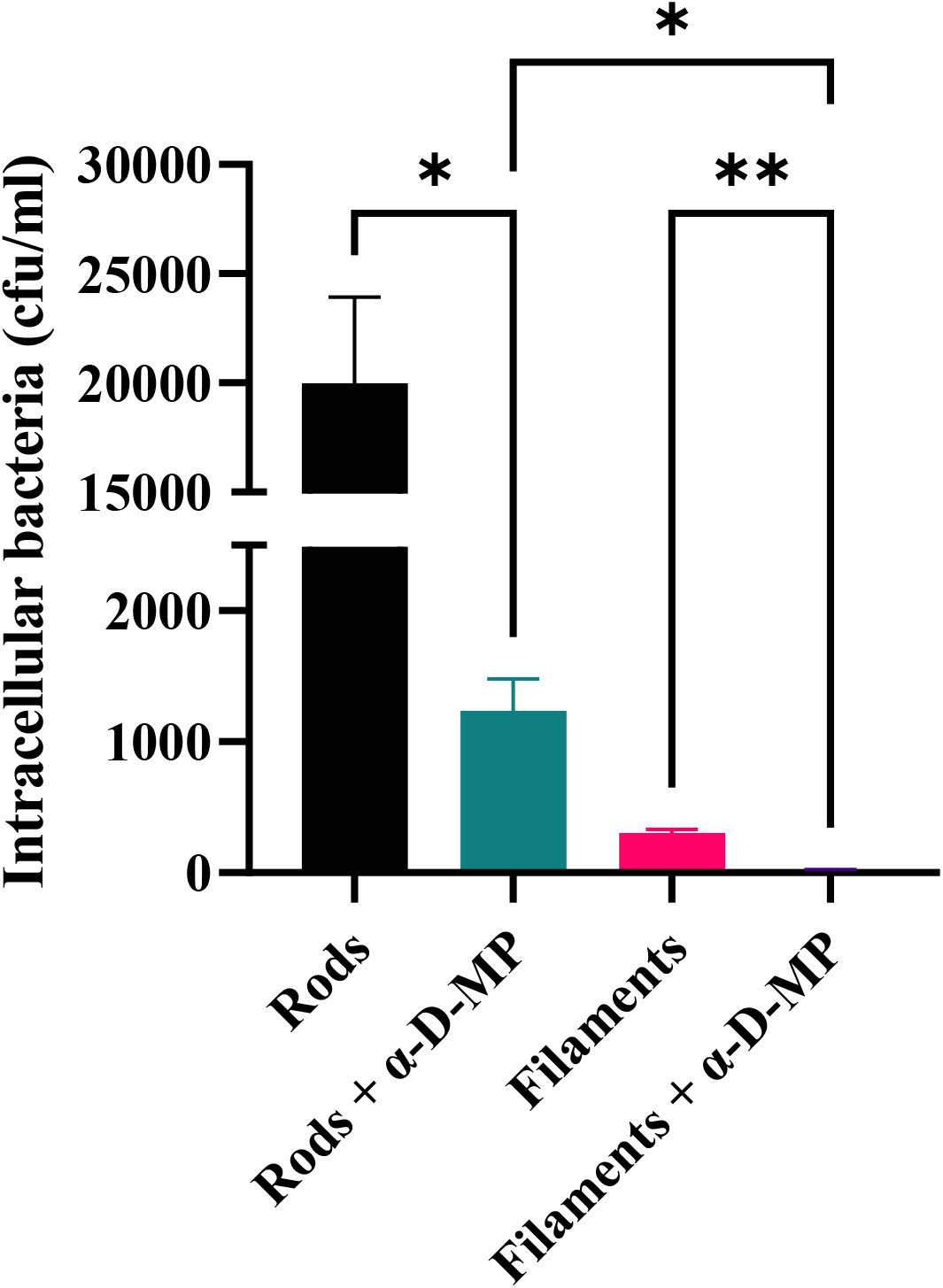
Blocking mannose binding with methyl α-D-mannopyranoside reduces THP-1 macrophage engulfment of UTI89 rods and filaments. *UTI89 was untreated (rods) or treated with 10 μg/ml cephalexin (filaments). Both populations were also treated with 3% methyl a-D-mannopyranoside (rods + α-D-MP and filaments + α-D-MP). THP-1 macrophages were infected (MOI 10) for 1 hour and intracellular bacterial loads were assessed 2 hours post infection in a gentamicin-protection assay. Data are the averages of 3 independent experiments with error bars representing the SEM. * indicates* p < *0.05*, ** p < *0.01, determined by one-way ANOVA with multiple comparisons*.

### The preference for macrophages to engulf rods more readily than filaments is abolished by deleting *fimH*

We show that the substantial decrease in engulfment of filaments relative to rods is not due to a decreased affinity of these cells for the mannosylated surface of macrophages. A primary mechanism of mannose-dependent macrophage internalisation is via type 1 fimbriae on the bacterial cell surface, where the tip-associated FimH protein binds mannosylated residues on the CD48 surface receptor of macrophages with high affinity [58]. Since α-D-MP blocks both UTI89 rods and filaments binding to macrophages to a similar degree, it would be expected that the deletion of *fimH* would give a similar result.

We confirmed that UTI89Δ*fimH* rods and cephalexin-induced filaments lacked functional type 1 fimbriae using a yeast agglutination assay. Exogenous expression of *fimH* from a plasmid restored the ability of both rods and filaments to agglutinate yeast indicating functional type 1 fimbriae expression could be complemented (Table S2 and accompanying text). Engulfment efficiency was determined by gentamicin-protection assays and yielded two unexpected observations. First, and in stark contrast to the mannose-binding inhibition assay (Fig 5), we found that both Δ*fimH* rods and Δ*fimH* filaments were engulfed to a much greater degree than wild-type rod and filament populations (Fig 6A); a 20-fold increase (*p* = 0.044) for rods and over 1000-fold increase (*p* = 0.049) for filaments compared to their wild-type counterparts. Furthermore, the increase in engulfment in the absence of *fimH* abolished the differing ability of macrophages to engulf filaments compared to rods (*p* > 0.05). Widefield fluorescence microscopy analysis of fully engulfed bacteria (bacteria completely enveloped by the macrophage membrane) confirmed that there was no significance difference (*p* > 0.05) in the engulfment of Δ*fimH* rods and Δ*fimH* filaments; an average 0.6% of the Δ*fimH* Rod population added were fully engulfed by each macrophage, compared to an average 0.4% of the Δ*fimH* Filament population (Fig 6B-D). While complementation with *fimH* (pGEN-*fimH*) did not reduce engulfment to wild-type levels for either rods or filaments (Fig S8), the yeast agglutination assay showed that functional type 1 fimbriae were restored in the complemented rods and filaments (Table S2), suggesting that the pathway of UTI89 engulfment was not solely through type 1 fimbrial binding to mannosylated THP-1 cell surfaces. Supporting this hypothesis, adding α-D-MP to Δ*fimH* rods and Δ*fimH* filaments also did not reduce engulfment to wild-type levels (Fig S9). Thus, the engulfment of the Δ*fimH* strain appears to be through a mannose-independent pathway and a size-independent pathway, suggesting that under certain conditions, as well as bacterial cell size and shape, additional FimH-dependent factors are influencing internalisation of filaments.

**Fig 6.**
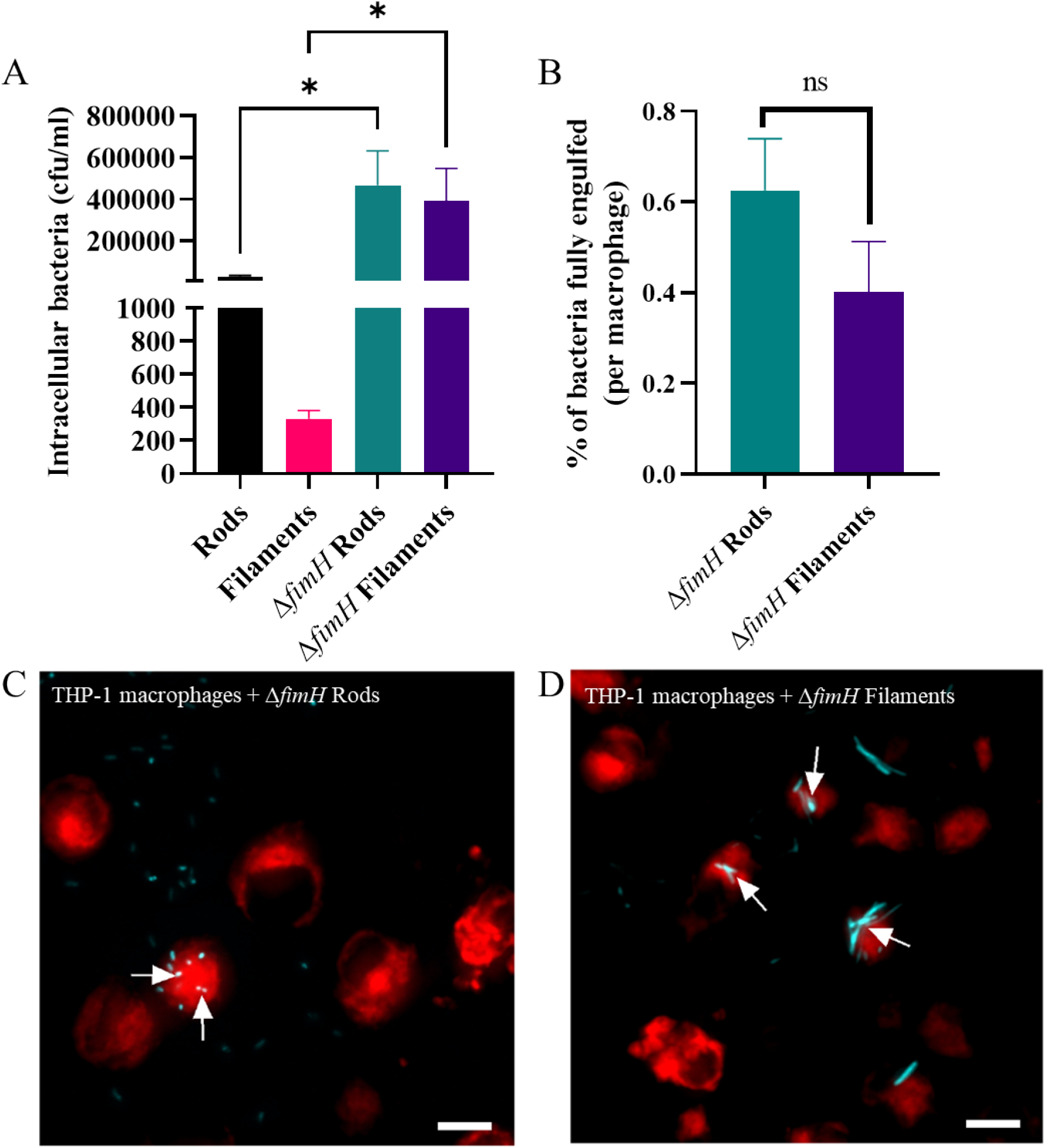
Deleting fimH abolishes the THP-1 macrophage preference to engulf rods more effectively than filaments. *UTI89, UTI89ΔfimH and UTI89ΔfimH/pGI5 (msfGFP) were untreated (rods or ΔfimH rods) or treated with 10 μg/ml cephalexin (filaments or ΔfimH filaments). (A) THP-1 macrophages were infected (MOI 10) for 1 hour and intracellular bacterial loads were assessed 2 -hours post infection in a gentamicin-protection assay. (B-D) Macrophages were infected with UTI89ΔfimH/pGI5 (msfGFP) rods (cyan) and UTI89ΔfimH/pGI5 filaments (cyan) at MOI 10, fixed after 60 minutes and stained with 1X CellMask™ Orange (red). Images were acquired using a DeltaVision Elite microscope with the 40X dry NA 0.60 objective. Internalisation of bacteria was determined using a FIJI macro on 3D image data. (B) Percentage of bacterial populations fully engulfed (number of bacteria fully engulfed/total number of bacteria counted) is normalised and presented as per macrophage to allow for comparisons despite variations in number of macrophages counted from microscopy images. (C-D) Representative images (maximum intensity projections) of infected macrophages with arrows indicates internalised ΔfimH rods (C) or ΔfimH filaments (D). Scale bar = 10 μm. Data are the averages of 2 independent experiments with error bars representing the SEM. * indicates* p < 0.05 *and ns indicates* p > *0.05, determined by one-way ANOVA with multiple comparisons or Welch’s t-test*.

### Bacterial growth environment contributes to macrophage engulfment dynamics

We have established that UTI89 cell size and shape both contribute to the effectiveness of macrophage engulfment under certain conditions, and that other factors such as FimH may contribute to engulfment efficiency. The environmental conditions under which bacteria are grown could influence these factors. Whilst *E. coli* forms filaments in response to certain antibiotics, UPEC filaments have also been observed in the urine of patients with UTIs who have not undergone antibiotic treatment [17]. Furthermore, UTI89 forms filaments inside human bladder cells *in vitro* when induced by the flow of concentrated urine [18, 28, 44]. We therefore examined the ability of THP-1 macrophages to engulf UTI89 isolated from an *in vitro* human bladder model which has not been exposed to antibiotic treatment. This replicates a heterogeneous bacterial population of varying cell lengths, from very short cells to very long, and such populations have been observed in UTIs of mice and humans [17, 44, 45, 59].

Bacterial populations (UTI89/pGI5 (msfGFP)) were isolated from the *in vitro* human bladder model and found to have a heterogeneous size distribution ranging from 0.7 μm – 235 μm, with ~ 86% classified as filamentous (cells > 4 μm in length) (Fig S10). Despite many filamentous bacteria in this population, macrophage engulfment of the mixed population of cells was higher than expected in the gentamicin-protection assay; equivalent to a rod population grown in LB (Fig 7A), and 100-fold higher than a cephalexin-treated filament population grown in LB. Fluorescence microscopy was used to quantify the proportion of rods and filaments being internalised by macrophages. Importantly, as for LB-derived cells, bladder rods are internalised more frequently than bladder filaments (0.65% vs. 0.24% of the bacterial population per macrophage respectively, *p* = 0.004, Fig 7B). Further classification of engulfed cells as either partial (incomplete enveloping of bacteria by macrophage membrane) or full (bacteria completely enveloped by the macrophage membrane) engulfment revealed that overall, a significantly higher proportion of filaments were partially (rather than fully) engulfed than their rod counterparts from both bladder-and LB-derived environments. It appears that the environment from which the filaments are derived contributes to the ability of macrophages to attempt engulfment, as significantly more bladder-derived filaments were partially engulfed compared to LB-derived filaments (*p* = 0.003, Fig 7C). Furthermore, there was no significant increase in partial engulfment of bladder-derived rods compared to LB-derived rods (*p* > 0.05), suggesting that this environmental contribution to engulfment also shows a size-dependency. Thus, bacterial cell length remains an important factor influencing successful macrophage engulfment (internalization) even when filaments are derived from different environments. Furthermore, our data indicates that specific environmental conditions may contribute to macrophage engulfment efficiency or dynamics, as macrophages attempt to engulf more filaments derived from a bladder infection model than those filaments grown in LB with cephalexin.

**Fig 7.**
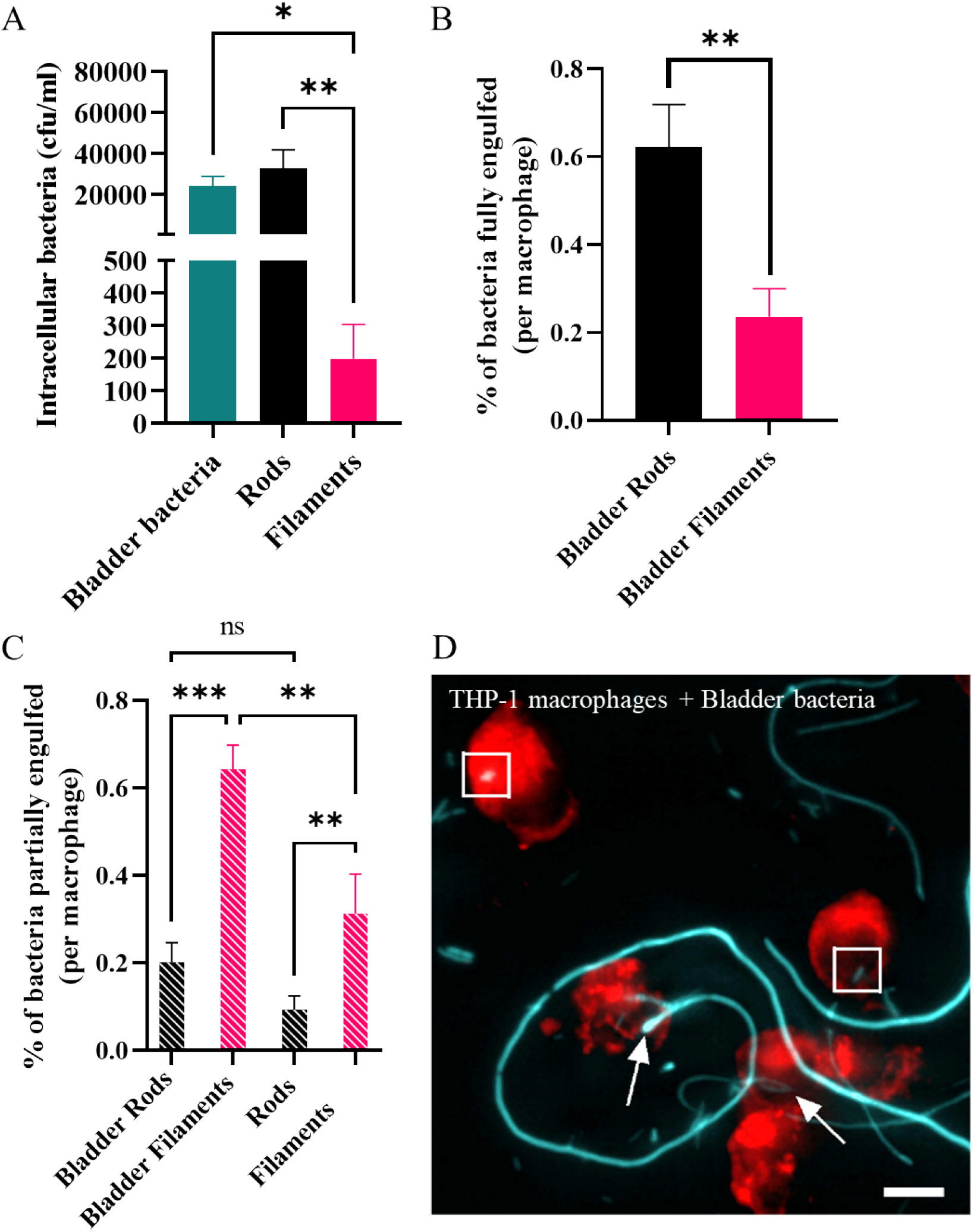
The growth environment of bacteria contributes to engulfment dynamics. *UTI89/pGI5 (msfGFP) were isolated from an in vitro human bladder model (bladder bacteria; ~86% filaments) or treated with 10 μg/ml cephalexin (filaments) or untreated (rods) and grown in LB medium. (A) THP-1 macrophages were infected (MOI 10) for 1 hour and intracellular bacterial loads were assessed 2-hours post infection in a gentamicin-protection assay. (B-D) Macrophages were infected with bladder bacteria (cyan) at MOI 10, fixed after 60 minutes and stained with 1X CellMask*™ *Orange (red). Images were acquired using a DeltaVision Elite microscope with the 40X dry NA 0.60 objective. Internalisation of bacteria was determined using a FIJI macro on 3D image data. (B) Percentage of bacterial populations fully engulfed (number of bacteria fully engulfed/total number of bacteria counted) is normalised and presented as per macrophage to allow for comparisons despite variations in number of macrophages counted from microscopy images. (C) Percentage of bacterial populations partially engulfed per macrophage as determined by microscopy presented as in B. (D) Representative image (maximum intensity projection) of infected macrophages with arrows indicating bacteria partially engulfed and boxes indicating bacteria fully engulfed. Scale bar = 10 μm. Data are the averages of 3 independent experiments with error bars representing the SEM. * indicates* p < *0.05*, ** p < *0.01 and ns indicates* p > *0.05, determined by one-way ANOVA with multiple comparisons*.

### UPEC filamentation in a mouse model and an *in vitro* human bladder model occurs independently of the SOS-induced filamentation genes, *sulA* and *ymfM*

One possibility to explain the increased attempts of macrophages to engulf human bladder cell-derived filaments, compared to filaments obtained in LB could relate to the mechanism by which filaments are formed in different environments. Previous *in vitro* studies have shown that cephalexin-induced filamentation by *E. coli* (strain SC1088), while inhibiting division by targeting PBP3, also induces the SOS response [60]. It has also been postulated that the SOS response, triggered by immune cell attack (oxidative damage), is required for UPEC infection in a mouse model of UTI [9, 61]. A major division inhibitor of the SOS response in *E. coli* is SulA, which is activated in response to DNA damage and interacts with FtsZ to inhibit the formation of Z rings and thus division [9, 62]. However, it has also been reported that UTI89Δ*sulA* was able to filament as efficiently as its wild type counterpart in an *in vitro* human bladder flow model of UTI [16, 18], suggesting that the intracellular infection-mediated UPEC filamentation occurs independently of SulA. Filamentation of *E. coli in vitro* has also been observed to occur in the absence of the SulA [16, 63–65].

We recently identified a e14 prophage gene, *ymfM*, that contributes to filamentation during the SOS response [65]. We therefore investigated if UTI89 still undergoes filamentation in the absence of both SulA and YmfM in a mouse cystitis model. Mice were infected with UTI89Δ*sulA*/pMAN01 (GFP-expressing plasmid), UTI89Δ*ymfM*/pMAN01, UTI89Δ*sulA*Δ*ymfM*/pMAN01, or wild type cells, UTI89/pMAN01. Filaments, which were morphologically indistinguishable from the wild-type (Fig 8A), were observed by confocal laser scanning microscopy on the surface of infected mouse bladders in the absence of *sulA* or *ymfM* (data not shown), as well as in the double mutant (Fig 8B), indicating that both these SOS-mediated filamentation genes are dispensable for UPEC filamentation under these conditions.

**Fig 8.**
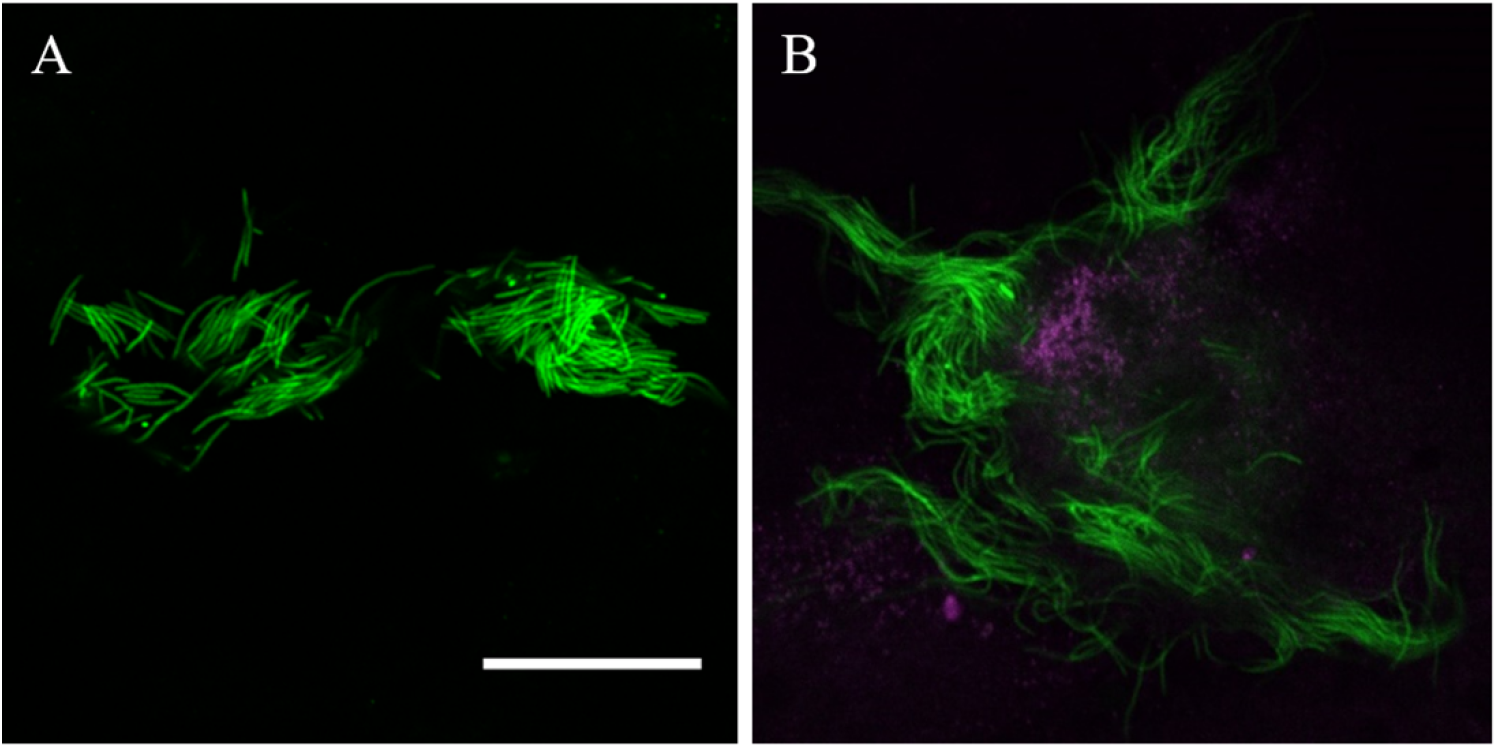
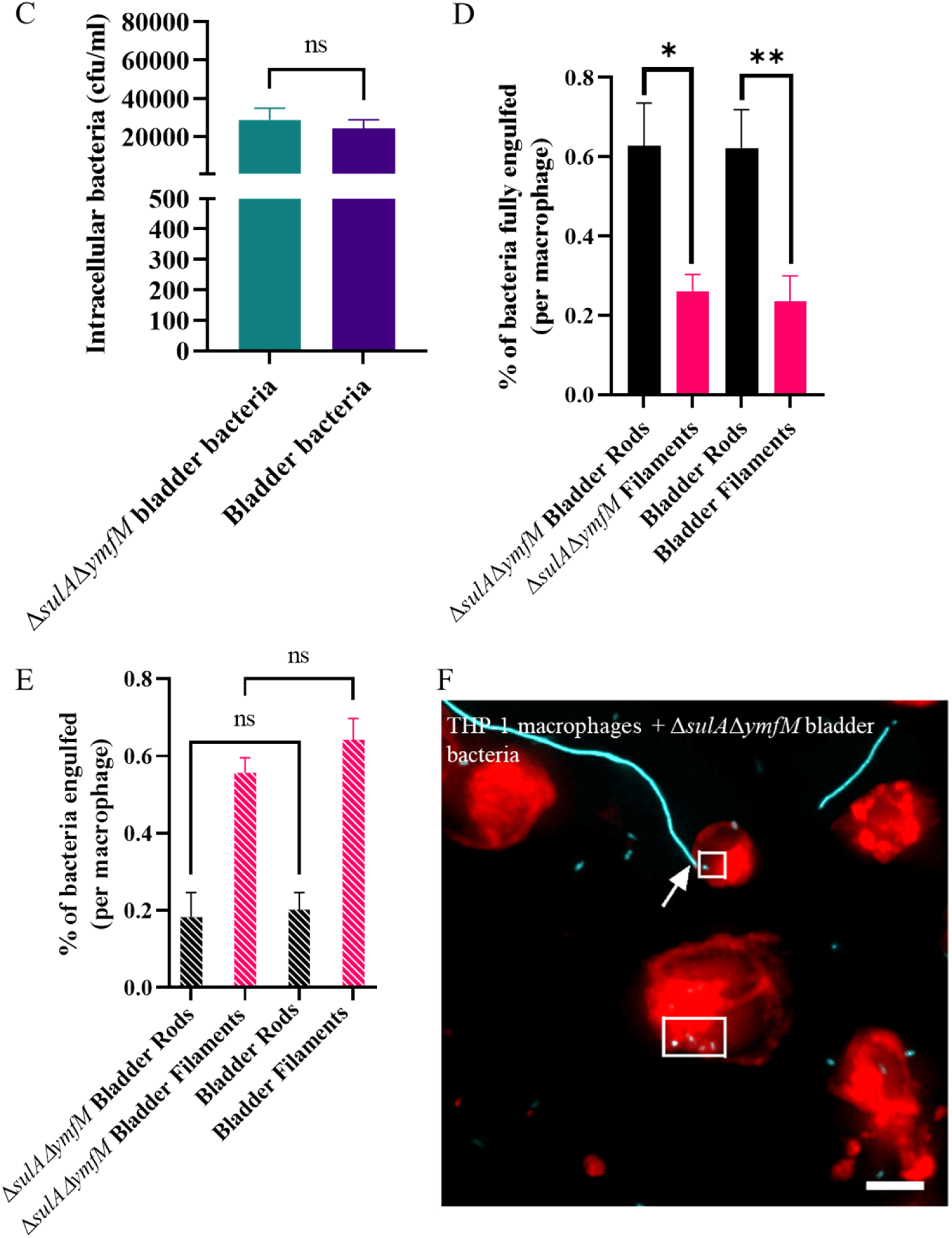
UTI89 mutants lacking sulA and ymfM retain the ability to form filaments and replicate engulfment dynamics of wild-type UTI89. *(A-B) Six hours post-infection, with (A) UTI89/pMAN01 (GFP) or (B) UTI89ΔsulAΔ*ymfM/*pMAN01 (GFP), C3H/HeN mouse bladders (magenta) were bisected and splayed on a silicone pad, fixed and imaged by an Olympus FV1000MPE microscope with a 20X objective (NA 0.75). Each bacterial strain was tested on a group of 4 mice with images representative from 1 mouse. Scale bar = 50 μm. (C-F) UTI89/pGI5 (msfGFP) or UTI89ΔsulAΔ*ymfM/*pGI5 (msfGFP) were isolated from an in vitro human bladder model (bladder bacteria or ΔsulAΔ*ymfM *bladder bacteria respectively). (C) THP-1 macrophages were infected (MOI 10) for 1 hour and intracellular bacterial loads were assessed 2-hours post infection in a gentamicin-protection assay. (D-F) Macrophages were infected with bladder bacteria or ΔsulAΔ*ymfM *bladder bacteria (cyan) at MOI 10, fixed after 60 minutes and stained with 1X CellMask*™ *Orange (red). Images were acquired using a DeltaVision Elite microscope with the 40X dry NA 0.60 objective. Internalisation of bacteria was determined using a FIJI macro on 3D image data. (D) Percentage of bacterial populations fully engulfed (number of bacteria fully engulfed/total number of bacteria counted) is normalised and presented as per macrophage to allow for comparisons despite variations in number of macrophages counted from microscopy images. (E) Percentage of bacterial populations partially engulfed per macrophage as determined by microscopy presented as in D. (F) Representative image (maximum intensity projection) of infected macrophages with arrows indicating bacteria partially engulfed and boxes indicating bacteria fully engulfed. Scale bar = 10 μm. Data are the averages of 3 independent experiments with error bars representing the SEM. * indicates* p < *0.05*, ** p < *0.01 and ns indicates* p > *0.05, determined by one-way ANOVA with multiple comparisons*.

To investigate the engulfment efficiency of filaments formed in the absence of *sulA* and *ymfM*, a population of heterogeneous lengths (UTI89Δ*sulA*Δ*ymfM*/pGI5 (msfGFP)) was derived from the *in vitro* human bladder model. Upon harvesting the bacteria from the flow cell model, phase-contrast microscopy showed that *UTI89ΔsulAΔymfM/pGI5* had high yields of highly filamentous bacteria (> 20 μm) present, indistinguishable from the wild-type UTI89/pGI5 (*UTI89ΔsulAΔymfM/pGI5;* data not shown, UTI89/pGI5 lengths in S10). This was complemented with flow cytometry, which showed similarly placed histogram peaks for both strains, indicating that both strains contained the same proportion of filamentous bacteria (data not shown). A gentamicin protection assay showed there was no difference in macrophage engulfment levels in the absence of both SulA and YmfM compared to UTI89/pGI5 (msfGFP) from the *in vitro* human bladder model (Fig. 8C). Fluorescence microscopy indicated that macrophage engulfment dynamics of both rods and filaments lacking both *sulA* and *ymfM* were comparable to their wild-type counterpart: there was still a preference (2.4-fold, *p* = 0.018) for internalisation of rods compared to filaments (Fig 9D and F), and macrophages partially engulfed Δ*sulAΔymfM* filaments grown in the *in vitro* bladder (0.56%) in similar proportions to UTI89 bladder filaments from the same model (0.64%) (Fig 9E). Together, these data show that UTI89 is still able to filament during *in vivo* and *in vitro* infection models in the complete absence of the two known SOS filamentation genes *ymfM* and *sulA*, and that macrophages interact with these filaments in a similar way to wild-type human bladder-derived filaments.

## Discussion

Bacterial filaments have historically been regarded as by-products of growth in ‘stressful’ or toxic environments. However, we are now beginning to understand that morphological plasticity may provide an advantage to many bacteria in a variety of environments. One such example is that UPEC filaments delay/inhibit engulfment by immune cells such as neutrophils and macrophages in a mouse model of UTI [8, 20]. Whilst it is assumed that this reduced engulfment is due to increased bacterial length, the exact advantage of filamentation has remained unclear. There is also an implicit assumption that all filaments are biologically, chemically, and structurally the same, and filament length or the environmental cues to induce filamentation are often overlooked as factors that may influence macrophage engulfment. We have begun to address these fundamental questions by examining the minimum cell length needed to significantly reduce engulfment by macrophages, and by systematically investigating biophysical parameters such as size and shape, and physiological parameters such as surface and environment, to understand what factors contribute to the filament survival advantage in the presence of human macrophages.

During infections bacteria are not inert or dead. Here we used live, viable populations of bacteria, more truly reflective of how they interact with immune cells. For the first time, to the best of our knowledge, this study identified conditions that produced viable, metabolically active and transient filaments of known lengths. This is critical to understanding how a changing morphology provides a survival advantage in the environment. In support of previous studies [8, 20, 21], we found that UPEC filaments (bacteria > 4 μm long) were engulfed significantly less than rods – up to 150-fold less - (Fig 1B-D) and in mixed populations rods were engulfed preferentially over filaments (Fig 2, Fig 7 and Fig 8). Further analysis found that UPEC populations, exposed to antibiotics, only needed as little as a doubling in average cell length to be less effectively engulfed, and the level of engulfment decreased exponentially as bacterial cell length increased (Fig 3). In simple terms, getting as long as possible in the environment could provide bacteria with an increased survival advantage. However, the energetic burden of switching from cell division to filament formation and back is unknown. Previous studies have shown links between central metabolism, cell division and cell size [66, 67], and proteomics analysis of *E. coli* filaments (induced by ampicillin treatment of persister cells) has demonstrated changes in carbon metabolism [68], but the direct link between filamentation and metabolism has not yet been fully investigated. Regardless of the energetic cost or benefit of filamentation, it is clear that bacterial cell length is a major contributor to macrophage engulfment efficiency.

Computational predictions of engulfment [25] and practical studies using inert particles [50] or bacterial samples [21, 22] have all demonstrated that successful engulfment of filamentous shapes, regardless of their size, relies on the phagocyte attaching to the pole of the target. In fact, the actual engulfment of *Legionella* filaments occurs at the same speed as rods but is delayed by the macrophage trying to reorient the filament to find the pole [21]. The bacterial populations from the *in vitro* bladder model used here included cells of various sizes with some filaments being over 250 μm long, over 10-fold longer than cephalexin-induced filaments. Investigating engulfment efficiency (comparing partially internalised and fully internalised bacteria) in this study revealed that macrophages partially internalised more filaments compared to rods, in both LB-derived and bladder-derived populations, with increased attempted engulfment of filaments from the *in vitro* bladder model. While not captured during the time frame of our assay, we hypothesize that this captured partial internalisation may eventually result in greater numbers of fully internalised filaments with engulfment being delayed by the requirement of finding the pole of the very long filaments, such as those in the bladder-derived population. A previous theoretical (computational) study showed that increasing sphere size would decrease engulfment by increasing the time required for membrane wrapping of the target [69]. Our work supports this hypothesis, we observed that as sphere volume increases (‘big spheres’), they are engulfed 67-fold less than the smaller spheres. Therefore, for UTI89 of different shapes, cell size remains an influence on effective engulfment.

While the size and shape of engulfment targets are closely linked, the shape of targets can additionally be described by aspect-ratio, curvature and circulatory. On examining how UTI89 cell shape might affect macrophage engulfment, we found that spherical cells are preferentially engulfed over rods, even when rods have a smaller volume. Spherical cells are also engulfed significantly more than filaments of approximately equal volume, indicating that UTI89 shape, as well as length and size, influence the ability of macrophages to engulf these bacteria (Fig 4A). This result builds on previous engulfment studies, using viable bacteria instead of inert particles. These previous studies found that engulfment relies on the circularity of nanoparticles [70], and that macrophage engulfment of spherical polystyrene particles is greater than that of high-aspect ratio shapes such as “worm-like” particles, of equal volumes [50]. Modes of entry of targets through a membrane have been predicted to differ based on the size, aspect ratio and curvature of particles, although the predicted mode of entry for high-aspect ratio and round tipped particles was noted to be different to that demonstrated practically for bacterial filaments [25]. Thus, our results both support and build on those described in previous studies and demonstrate an interplay between size and shape of bacteria that affects macrophage engulfment.

Whilst spheres produced by mecillinam treatment were viable in this study, it was noted that viability was reduced for mecillinam/cephalexin treated spheres compared to untreated rods. This could be due to a number of reasons: cephalexin and mecillinam are known to have a synergistic effect on *E. coli* viability [71], with links to the bacterial metabolite bulgecin [72] as well as other metabolic changes induced by mecillinam alone [73]. Treatment of *E. coli* with sub-lethal mecillinam has also been demonstrated to induce cell wall changes, making it more isotropic and having altered chirality [74]. Although the chemical composition of the cell wall has been reported to be unaltered after mecillinam treatment [74], the addition of cephalexin used in this study may contribute further to cell wall changes. It has been reported that cephalexin increases the amount of structural change in the cell wall when used in addition to another penicillin antibiotic, ampicillin [75]. So, while increasing the size of spheres did reduce engulfment, further investigation on the role of surface composition should be performed to determine whether the macrophage engulfment changes were due to alterations in cell wall composition. Genetic manipulation of *E. coli* through deletion of the rod-shape determining protein MreB [76] could be informative in investigating macrophage engulfment of round cells in the absence of antibiotic treatments.

The role of Type 1 fimbriae and the adhesin FimH in both establishing infection and engulfment through binding to mannosylated glycoproteins on mammalian cells has been well established for rod-shaped *E. coli* [77]. A previous study deleted the entire *fim* operon from a K-12 strain of *E. coli* and found that these bacteria do not get engulfed by RAW 264.7 mouse macrophages [78], supporting the theory that fimbriae are essential for phagocytosis of *E. coli*. We found indirect support for the importance of fimbriae in engulfment by blocking the ability of UTI89 to bind to mannose and observing a reduction in engulfment by macrophages (Fig 5). The equal drop in engulfment of rods and filaments indicates that there is no difference in their ability to interact with the mannosylated host cell surface, and that size/shape of bacteria is the primary driver of engulfment efficiency and not an inability of filaments to bind to the mannosylated surface of macrophages. However, low-level engulfment of both rods and filaments was still observed even after the blocking of available mannose-binding sites, suggesting that binding to the mannosylated host cell surface may not be the only pathway for engulfment. Macrophages have numerous receptors on their cell surface, performing a variety of functions that contribute to phagocytosis: some initiating signalling pathways to induce phagocytosis directly while others work in conjunction with these phagocytic receptors to detect molecular patterns of pathogenic invaders [79]. The vast numbers and types of receptors capable of phagocytosing different targets through various pathways [79] coupled with the range of glycans and glycolipids on the UPEC cell surface [56, 80], and the ability of UPEC to switch expression of its surface structures on/off [81, 82], supports this hypothesis that alternate engulfment pathways do exist.

Unexpectedly, deleting *fimH* in UTI89 increased engulfment of both rods and filaments and abolished the observed size/shape dependency. Thus, the engulfment of the Δ*fimH* strain appears to be through a mannose-independent pathway and a size/shape-independent pathway. This suggests that under certain conditions the biophysical aspects of bacteria are not the only major contributors to engulfment and other factors such as bacterial cell surface are playing a role. Reduction in T cell activation and dendritic cell migration has been reported as a consequence of FimH binding to CD14 on the surface of dendritic cells, implying that FimH may be involved in immune cell evasion [83]. One challenging aspect of this study is understanding the cellular changes caused by changing cell size, either directly due to increased length/volume, or indirectly in response to filament induction methods, and the role these play in macrophage engulfment efficiency. It cannot be ruled out that the loss of size/shape dependency for macrophage engulfment is due to the combined effect of the *fimH* deletion and the use of cephalexin targeting the bacterial cell wall. Expression of type 1 fimbriae is phase variable [81] and, though cells were selected for at the population level in this study (by static growth), the amount of expression or whether individual bacteria were expressing fimbriae was not determined. Deletion of *fimH* has been shown to result in a large reduction of surface-associated type 1 fimbriae [84], therefore assessing the proportion of fimbriated bacteria in both wild-type and Δ*fimH* rods and filaments would offer valuable insights into fimbriation during filamentation. Whilst it is currently unknown what other surface structures, pathways, or global cellular responses could be affected by deleting *fimH*, further investigations are highly warranted.

Macrophage engulfment of bacterial populations isolated from an *in vitro* bladder model show that they attempt to engulf more filaments compared to filaments grown in defined nutrient medium. This environmental effect during filament development in the *in vitro* bladder model is currently unknown, however could be due to complement-coating or opsonization which have previously been shown to enhance the engulfment of certain pathogens [85, 86]. It has also been demonstrated that UPEC undergoes filamentation when exposed to urine in both the presence and absence of host bladder cells [18, 28], which raises the possibility that unknown component(s) of urine could influence macrophage engulfment. UTI89 planktonic bacteria cultured in human urine can be devoid of type 1 fimbriae, and switch to expressing type 1 fimbriae when in sessile populations [87]. A lack of functional type 1 fimbriae was identified in this study to increase macrophage engulfment and thus could be responsible for the increased engulfment attempts of filaments from the bladder model.

We found that UTI89 was still able to filament when exposed to urine in the complete absence of the two known SOS filamentation genes *ymfM* and *sulA* in both an *in vivo* mouse UTI model and an *in vitro* human bladder cell model. Previously, deletion of *sulA* was reported to abolish filamentation in an *in vivo* mouse model of UTI [20], whilst an *in vitro* flow cell model of UTI still resulted in filamentation in UTI89Δ*sulA* [16, 18]. These differences may be due to differences in the timing of observation of filamentation, or differences in bacterial strain construction in these studies. Our results support the proposal that known components of the SOS response are dispensable for UPEC filamentation during the establishment of UTIs. Furthermore, we demonstrate that the increased ability of macrophages to partially engulf human bladder-derived filaments is unchanged in their absence. Our data is in agreement with a previous study where both wild-type and Δ*sulA* UTI89 filaments (induced by mitomycin C treatment) were readily engulfed by human and mouse polymorphonuclear leukocytes [20]. Overall, there are likely many unknown factors that induce filamentation under different environmental conditions, and how these influence the dynamics and efficiency of macrophage engulfment requires future investigations.

In conclusion, the data presented here contributes to our understanding of why bacteria filament during UTIs but also highlights the pitfalls of making general conclusions when discussing bacterial filamentation. There are many factors that influence the interactions between the bacteria and host, and it is becoming increasingly clear that these interactions are incredibly complex and warrant further study. Some of the bacterial factors that affect host macrophage engulfment (and thereby infection responses) were investigated here, revealing that for *E. coli* UTI89, size, shape, surface and growth environment can cause variations in the engulfment effectiveness by human THP-1 macrophages. Furthermore, the long-known SOS response of bacteria still has many unknown aspects and the novel observation of engulfment differences of filaments formed by antibiotics in LB and within bladder epithelial cells in urine open new research directions for future exploration. Improving our knowledge of bacterial filaments will allow a deeper understanding of the process of infection, how pathogens interact with our immune response, and ultimately uncover novel avenues for non-antibiotic treatments which are urgently needed to treat resistant pathogens.

## Methods

### Table S3 and Table S4 contain lists of bacterial strains, plasmids, and primers

#### Conditions to establish filamentation

##### Growth of rods and antibiotic-induced filamentous bacteria

All bacterial strains were grown statically at 37°C unless otherwise stated. Twenty-five ml of LB was inoculated from an overnight culture to a starting OD_600_ = 0.05 and incubated statically until early exponential phase (OD_600_ ~ 0.2). One culture then acted as a control with no antibiotic addition, whilst the remaining two had 2.5 μg/ml (LEX rods) and 10 μg/ml (LEX filaments) of cephalexin added. Cultures were grown until OD_600_ of control reached 0.7-0.8, before cells were centrifuged at 3500 *g* for 5 minutes and resuspended in PBS to CFU/ml of 2×10^7^. UTI89/pGI5 (msfGFP), *UTI89*Δ*fimH* and UTI89Δ*fimH*/pGI5 (msfGFP) was grown the same way with the addition of 100 μg/ml spectinomycin for selection of the pGI5 (msfGFP) plasmid in LB.

For mixed populations rods and LEX filaments were grown as described above with the following addition. After final centrifugation cultures were resuspended in PBS to a final CFU/ml of approximately 2×10^7^ of which 24% were rods (approximately 0.53 x 10^7^ CFU/ml) and 76% were filaments (approximately 1.6 x 10^7^ CFU/ml).

For ciprofloxacin-induced filaments UTI89 was grown the same way as described for cephalexin treatment with the following additions. After growth to early exponential phase one culture then acted as a control with no antibiotic addition, whilst the remaining two had 3.75 ng/ml (CIP rods) and 15 ng/ml of ciprofloxacin (CIP filaments) added. Cultures were grown until OD_600_ of rods reached 0.7-0.8, before centrifuged at 3500 *g* for 5 minutes and diluted to an OD_600_ of 0.1 in fresh LB and grown for 1.5 hours in the presence of the same concentrations of ciprofloxacin. Cultures were centrifuged at 3500 *g* for 5 minutes and resuspended in PBS to CFU/ml of 2×10^7^.

##### Expression of *ftsZ-yfp*

First, 5 ml of LB with ampicillin (100 μg/ml) and 0.2% (v/v) glucose solution (to repress expression of *ftsZ-yfp* from pLau80) was inoculated with a single colony of UTI89/pLau80 or UTI89/pG15/pLau80 and allowed to grow overnight at 37°C. The overnight culture was diluted in 20 ml LB with ampicillin (100 μg/ml), spectinomycin (100 μg/ml) and 0.2% (v/v) glucose solution so that the OD_600_ = 0.05. Cultures were grown at 37°C for 2 hours before being centrifuged at 3500 *g* and washed twice with fresh LB media to remove glucose. Cultures were diluted to OD_600_ = 0.1 in 20 ml LB with ampicillin (100 μg/ml), spectinomycin (100 μg/ml) and either 0.2% (v/v) glucose solution (repression of pLau80) or 0.2% (v/v) arabinose solution (expression of pLau80). Cultures were grown at 37°C for 2 hours and then cultures were centrifuged at 3500 *g* for 5 minutes and resuspended in PBS to CFU/ml of 2×10^7^.

##### Growing filaments of different lengths

Cultures were grown as previously described for antibiotic induced filaments and expression of *ftsZ* with the following additions. Cultures were grown to different ODs before induction of filamentation through addition of cephalexin or 0.2% (v/v) arabinose. They were collected at the same end point, an OD_600_ equivalent to an untreated culture (0.7-0.8). Cultures ended up being grown at 37°C in the presence of 10 μg/ml of cephalexin for 1 hour 30 min, 1 hour 15 min, 1 hour, and 45 min, or in the presence of 0.2% (v/v) arabinose solution for 2 hours 15 min, 1 hour and 30 min. Cultures were centrifuged at 3500 *g* for 5 min and resuspended in PBS to CFU/ml of 2×10^7^.

##### Growing bacteria in different shapes

Cultures were grown as previously described for antibiotic induced filaments with the following additions. Cultures were grown at 37°C for 1 hour 15 min to an OD_600_ ~ 0.2 (early exponential phase). Cultures had 10 μg/ml mecillinam only or 10 μg/ml mecillinam and 10 μg/ml cephalexin added and were grown at 37°C for 1 hour 30 min before they were centrifuged at 3500 *g* for 5 minutes and diluted to an OD_600_ of 0.1 and grown for 1 hour 30 min in the presence of the same concentrations of antibiotics. Cultures were centrifuged at 3500 *g* for 5 min and resuspended in equal volumes of PBS.

##### Quantification of length and volume

Bacteria were fixed with 3.7% formaldehyde (v/v) for 1 hour at room temperature prior to measurement. Lengths of bacteria were measured through phase contrast microscopy on the Zeis Axioplan2 with accompanying software. Volumes of bacteria were measured by the Multisizer M4e Coulter Counter Analyzer (Beckman). Samples fixed in formaldehyde solution were diluted to 1% (v/v) in filtered IsoFlow Sheath Fluid (Beckman). Of this, 200 μl was run through a 50 μm aperture tube, and data were collected over 400 bins ranging from 0.5 μm^3^ to 100 μm^3^ were measured.

##### Growth and culture of THP-1 monocytes

The human monocytic cell line used was THP-1 (ATCC^®^ TIB-202™), which was derived from an acute monocytic leukaemia patient. The THP-1 cells were cultured at 37°C, 5% CO_2_ in RPMI-1640 medium containing 10% (v/v) FBS (fetal bovine serum, heat inactivated) and 1% (v/v) GlutaMAX™. Approximately 2×10^5^ THP-1 cells/ml were added to a 24 well plate (1ml per well) and were stimulated to form macrophages with 30 ng/ml of PMA (Phorbol 12-myristate 13-acetate) for 48 hours. THP-1 cells were washed and resuspended in fresh RPMI-1640 medium containing 10% (v/v) FBS and 1% (v/v) GlutaMAX™ (RPMI culture media) and incubated at 37°C, 5% CO_2_ for 24 hours before infection with bacteria.

##### Gentamicin-protection assay

The bacterial cultures were added to macrophages at an MOI of 10 (MOI 10) per well (approximately 2×10^6^ viable bacteria per well determined by CFU/ml). Plates were centrifuged at 1000 *g* for 5 minutes and incubated for 1 hour at 37°C and 5% CO_2_. Cells were then treated with 200 μg/ml gentamicin for 1 hour at 37°C and 5% CO_2_. Cells were washed twice with PBS and lysed with 0.1% (v/v) Triton X-100 for 15 min, serially diluted in PBS, plated on LB agar and incubated at 37°C for 16-18 hours. Colonies were then counted, and CFU/ml was calculated. This was done in technical triplicate, with at least 3 biological replicates.

For blocking with methyl α-D-mannopyranoside the gentamicin-protection assay was performed as described above with the following additions. Before the addition of bacteria, RPMI culture media was supplemented with 3% (w/v) methyl α-D-mannopyranoside (α-D-MP) [58]. The bacterial cultures (also in 3% (w/v) α-D-MP) were added to macrophages at an MOI 10 per well (approximately 2×10^6^ bacteria per well).

For microscopy analysis macrophages were stimulated the same way and were infected with the MOI 10 for 1 hour. However, CellCarrier-96 well Black plates were used for the assay with 1×10^5^ THP-1 cells/ml added (200 μl per well). After the initial 1-hour incubation at 37°C in 5% CO_2_ macrophages were stained with a 1X CellMask™ Orange Plasma membrane Stain at 37°C for 10 minutes in the dark, washed twice with PBS and fixed by adding 3.7% (v/v) formaldehyde at 37°C for 20 minutes in the dark.

##### Imaging

Fixed and live bacteria were viewed using phase contrast and widefield fluorescence microscopy with a Zeiss Axioplan 2 fluorescence microscope and a 100X oil immersion NA 1.4 Plan Apochromat objective lens. The light source was a 100 W high pressure mercury lamp that passed through filter blocks for observing GFP (Filter set 09; 450-490 nm BP excitation, 515 nm LP barrier filter) and propidium iodide (Filter set 15; 546/12 nm BP excitation, 590 nm LP barrier filter). Images were taken using a Zeiss AxioCam MRm camera and analysed using AxioVision software version 4.8 (Zeiss). For widefield fluorescence microscopy of UTI89/pGI5 (msfGFP), the exposure time was maintained at 200 ms. For imaging membrane integrity staining of GFP and propidium iodide the exposure times were maintained at 100 ms and 200 ms respectively.

Three-dimensional z-stack acquisition was performed using a DeltaVision Elite widefield fluorescence microscope (GE Healthcare) using a 40X objective (OLYMPUS LUCPLANFLN 40X dry objective NA 0.60). The filter sets used were GFP (464-492 nm excitation) and TRITC (531-565 nm excitation). Each stack acquired consisted of 25-30 optical slices with intervals of 1.45 μm (this was optimised for subsequent deconvolution microscopy). Stacks were deconvolved using the softWoRx Enhanced Ratio deconvolution method (GE Healthcare).

##### Digital analysis of microscopy images of engulfment

Analysis of deconvolved images was done using FIJI software utilising both available plugins and custom macros. Thresholding and masking to identify the macrophages was based on membrane staining with 1X CellMask™ Orange. Bacteria were identified by msfGFP expression from the pGI5 plasmid. Nearest neighbour distance calculations were used to identify how close bacteria were to macrophages including bacteria within the membrane of a macrophage. For rods, a fully internalised bacterium was defined as having a value of > 1.0. For filaments, a fully internalised filament was assigned two values for the closest and furthest aspect of the bacteria and if both were negative this correlated to a fully internalised filament, while if only one was negative this indicated partial engulfment. The total number of macrophages in an image stack were counted, and the total bacteria per image stack to enable analysis and quantification of population percentages. A minimum of 4 images were analysed per time point. A minimum of 2 biological replicates were performed. Data were measured per image, with an average calculated from the replicate images. Images were not taken by a blinded investigator, but the software defined the measurements.

##### *In vitro* bladder infection model

The set-up was based on a previously established approach [18, 44, 45], with minor modifications. Briefly, on day one, flow chambers (IBIDI μ-Slides I^0.2^ Luer, Cat#: 80166) were seeded with PD07i epithelial bladder cells at a concentration of ~ 3×10^6^ cells in EpiLife Medium (Gibco, #MEPI500CA) supplemented with growth supplements and antibiotics (HKGS, #S0015, and 100μg ml^-1^ Pen/Strep, SIGMA #P4333). Seeded channels were left overnight to allow cells to adhere and multiply into a confluent layer. The next day, flow channels were connected to New Era pumps via tubing and 20 ml disposable syringes. Flow (15 μl min^-1^) of fresh EpiLife (supplemented with HKGS) without antibiotics was maintained for 18-20 hours. On day three, to induce infection, bladder cells were exposed to bacterial cultures (i.e., UTI89 grown statically at 37 °C overnight) at a concentration of OD_600_ 0.2 for 15 min at a flow rate of 15 μl min^-1^. Following this step, the media was changed back to EpiLife (supplemented with HKGS), after an initial flow of 100 μl min^-1^ to flush out the excess bacteria, and flowed for an additional 9 hours to allow bacteria to adhere to and invade the epithelial bladder cells. This step was followed by flow (15 μl min^-1^) of EpiLife (supplemented with HKGS) in the presence of 100μg/ml gentamycin for 20 hours to allow for the formation of intracellular bacterial communities (IBCs), as well as to remove any lingering extracellular bacteria. Following these 20 hours, the media was changed to sterile filtered human urine (with pH between 5.12-5.83 and Urine Specific Gravity of at least 1.024 g mL^-1^, [44]) with flow (15 μl min^-1^) for 18 - 20 hours to induce filamentation and dispersal of the bacteria from the bladder cells. Filaments were collected from the back-opening of the flow channels.

##### Mouse cystitis model

The model is based on Hung, Dodson (88). Deletion strains were constructed in UTI89 by the well-established lambda Red recombinase method [89]. A total of 4 strains were constructed - UTI89/pMAN01, UTI89Δ*sulA*/pMAN01, UTI89Δ*ymfM*/pMAN01, UTI89Δ*sulA*Δ*ymfM*/pMAN01. After 1 week of acclimatization, 8-9-week-old female C3H/HeN mice (Janvier Labs, France) were anaesthetized followed by inoculated by transurethral catheterization with approx. 1-2 x 10^7^ cfu of either strain diluted in 50 μl PBS (as determined by OD_600_). After 6 hours of infection, the mice were euthanized, and bladders were collected and bisected. Each sample was then splayed on a silicone pad and fixed in 3% (v/v) paraformaldehyde followed by confocal laser scanning microscopy using an Olympus FV1000MPE microscope with a UPLSAPO 20X/0.75NA objective and Olympus FV10-ASW software. Each strain was tested on a group of 4 mice.

##### Statistical analysis

*P*-values were determined by one-way ANOVA with multiple comparisons (for 3 or more conditions), or t-test (for less than 3 conditions) through GraphPad Prism software. Tukey, Šidák or Welch’s tests were used for multiple comparisons, and Tukey or Welch’s test for t-tests on less than 3 conditions, depending on the test for equal variance. Power analysis was performed on raw data to confirm the sample size required (α = 0.05 and desired power = 0.8), following this data was normalised if required.

##### Ethics statement

The mice experiments were conducted according to the national guidelines by the National Danish Animal Care Committee. The protocol was approved by the Danish Veterinary and Food Administration under the Ministry of Food, Agriculture and Fisheries with ID number 2015-15-0201-00480. Human urine collection was approved by the UTS Human Research Ethics Committee (HREC no. 2014000452).

## Acknowledgements

The authors acknowledge the use of the equipment: the Zeiss Axioplan2 fluorescence microscope with AxioVision version 4.8 software (Zeiss), Nikon TiE2 N-STORM and the DeltaVision Elite widefield fluorescence microscope and softWoRx software (GE Healthcare) in the Microbial Imaging Facility at the Australian Institute for Microbiology and Infection (AIMI) in the Faculty of Science, the University of Technology Sydney. We would like to thank Louise Cole and Chris Evenhuis for their scientific input and/or technical assistance. Bill Söderström is supported by a Discovery Project grant from the Australian Research Council (DP220101143). Finally, to Nilesh Bokil who assisted in starting work with macrophages. This work was supported by Australian Government RTP scholarship.

## Author Contributions

**Elizabeth Peterson:** Conceptualization, Methodology, Formal analysis, Investigation, Writing-Original Draft preparation, Visualization; **Bill Söderström:** Investigation, Writing – Review and editing; **Nienke Prins:** Methodology, Investigation; **Giang H.B. Le:** Investigation, **Lauren E. Hartley-Tassell:** Methodology, Investigation; **Chris Evenhuis:** Software; **Rasmus Birkholm Grønnemose:** Investigation, Writing – Review and editing; **Thomas Emil Andersen:** Supervision, Writing – Review and editing; **Jakob Møller-Jensen:** Supervision, Writing – Review and editing; **Gregory Iosifidis:** Methodology; **Iain G. Duggin:** Supervision, Writing – Review and editing; **Bernadette Saunders:** Supervision, Writing – Review and editing; **Elizabeth J. Harry;** Conceptualization, Funding Acquisition, Resources, Supervision, Writing – Review and editing; **Amy L. Bottomley:** Conceptualization, Investigation, Project Administration, Supervision, Investigation, Writing-Original Draft preparation.

